# Origami-based growing tube model for reproducing shell shapes

**DOI:** 10.1101/2023.07.19.549674

**Authors:** Maho Ueda, Nozomi Fukunaga, Noa Yamashita, Yuki Yokoyama, Hideaki Kida, Keisuke Matsuda

## Abstract

Mathematical models for shell morphology have been well-studied in theoretical morphology. The growing tube model proposed by Takashi Okamoto was developed to reproduce heteromorph ammonoids and can generate various shells by changing three parameters: curvature, torsion, and enlarging ratio. Previous studies in theoretical morphology have employed computers to visualize mathematical models. However, computers are not accessible to everyone, and merely observing the graphics of the reproduced shells does not deepen understanding of the mathematical model. Therefore, in this study, we considered using origami, which is inexpensive and readily available, to make users understand mathematical models through hands-on folding. First, we simplified the crease pattern of seashells, originally devised by an origami artist Tomoko Fuse, to create a basic crease pattern. We then identified the crease pattern elements corresponding to the three parameters in the growing tube model and the aperture-apex position determiner. Based on them, we reproduced various shell shapes, including a heteromorph-ammonoid-like shape, with origami. Finally, we investigated the constraints between parameters in the origami-growing tube model. The origami-growing tube model is expected to help disseminate mathematical models and promote theoretical biology.

## Introduction

Shells have not only attracted people but have also been the subject of scientific research. Shells are created by various animals, especially mollusks, and many shells have a spiral or coiled form.

In paleontology, especially in theoretical morphology, researchers have attempted to reproduce shell shapes using mathematical models based on the regularity of their spiral forms and the way of growth. Once a mathematical model that can approximate forms is developed, various shell forms can be generated by changing the model’s parameters, which generates the spectrum of theoretically possible forms (including forms that do not exist in reality). It allows putting each morphological type in a conceptual framework. In the spectrum, actual species are not randomly distributed. Investigating the distribution helps us consider the shape’s functional and developmental restriction, finally leading to understanding the morphogenesis mechanisms.

The shells of a mollusk, nautilus, or ammonite are made up of beautiful, regular spirals called “equiangular” or “logarithmic” spirals. The term “equiangular” is derived from the fact that the angle between the radius and the tangential line is constant at any point in the curve. The “logarithmic” is derived from the fact that when the spiral is expressed by polar coordinates, a logarithmic (or exponential) function appears (the following equation).

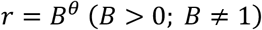

The equation has only one parameter *B*. Therefore, shells coiled on a flat surface can be reproduced by measuring *r* and θ at two points. Historically, Moseley proposed a geometric model for the shell coiling based on the equiangular spiral (Moseley 1838). Thompson reviewed this and other early studies on the coiling of mollusk and foraminifera (Thompson and Thompson 1942). After that, Fukutomi pointed out that the model is applicable not only to gastropods and nautiloids but also to bivalves (Fukutomi 1953). These data confirmed the general applicability of logarithmic spirals.

Based on logarithmic spirals, American paleontologist David M. Raup developed Raup’s model, which can be regarded as a logarithmic spiral with height displacement (Raup 1966). The model includes four parameters: *W* (whorl expansion rate), *D* (distance between the coiling axis and generating curve), *T* (whorl transition rate), and *S* (shape of the generating curve). The generating curve *S* is often equivalent to the outline of the growing edge of the shell. *W, D*, and *T* can be measured as simple ratios between two parts. So, by measuring the length with actual shells, the model’s parameters can be assessed. By varying the parameters, many coiling shell shapes can be reproduced, including most gastropods, cephalopods, and bivalves.

These models employ fixed axes, regardless of rectangular or polar coordinates, but any fixed coordinate system is no more than an artificial framework imposed upon the coiling pattern. For living organisms, at least, fixed coordinates are irrelevant to the fundamental mechanism of coiling.

Okamoto proposed a growing tube model that employs differential geometry (Okamoto 1988). The model can describe any coiled shell with a circular cross-section, with three differential parameters: *E* (radius enlarging ratio), *C* (standardized curvature), and *T* (standardized torsion). The equations in the models are:

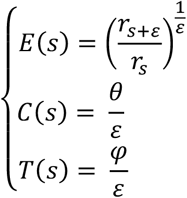

where each character is shown in Figure 1a. The calculation of each parameter includes 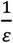 for normalization. This model was initially developed to explain the heteromorph ammonoids that look too complicated to explain how they are formed. The growing tube model makes the complicated shell form of heteromorph ammonoids considered simply as an integration of ad hoc accretional growth of the aperture without defining any coordinate system. This model can also generate most shell shapes by changing the parameters (Figure 1b).

**Figure 1:**
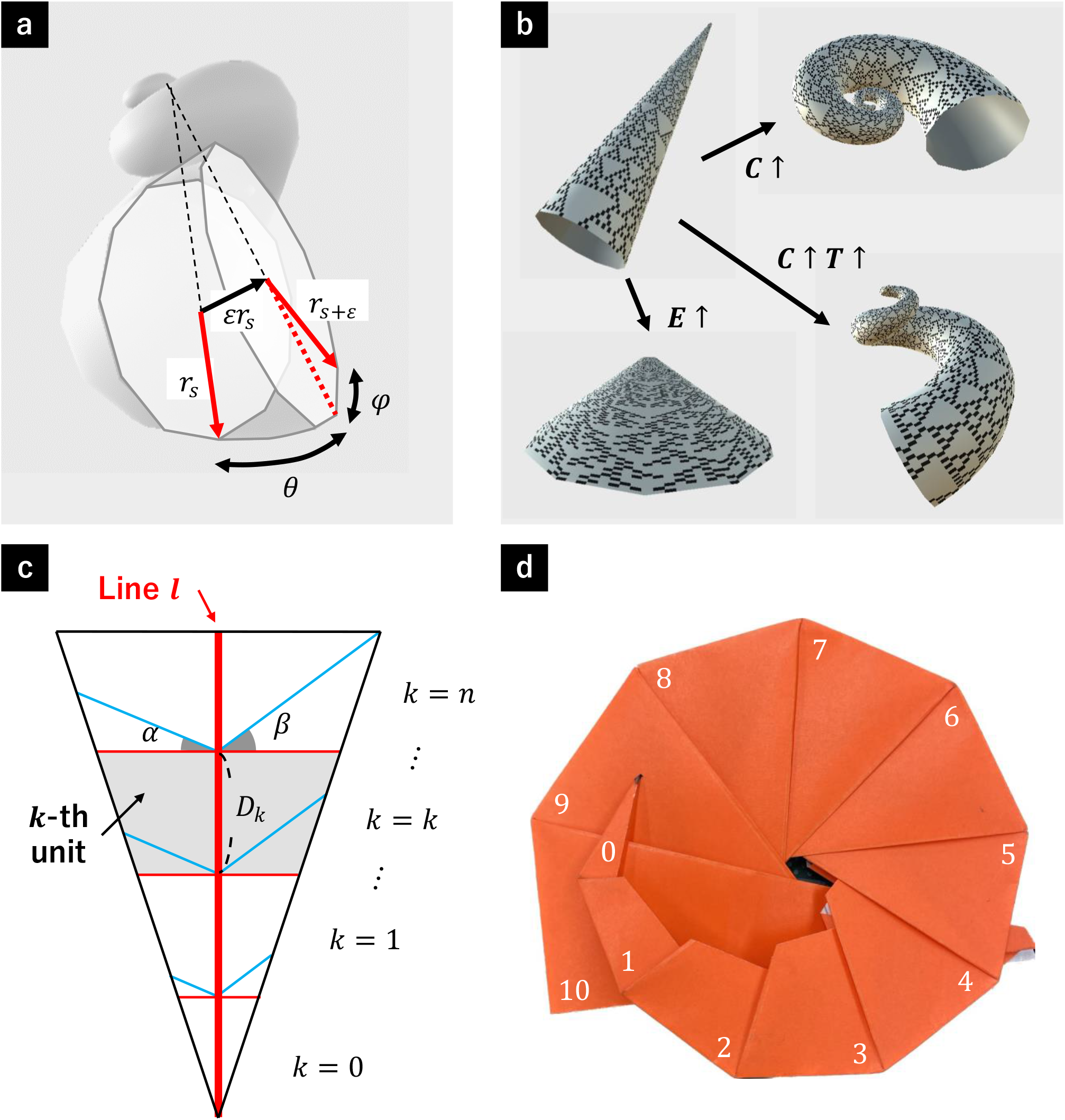
Basic crease pattern for the origami-growing tube model. (**a**) The relationships between the current stage *s* and the next stage *s* + ε in the growing tube model. *rs* denotes the radius of the aperture at stage *s*. The red and black arrows show the normal and tangent vector directions of the Frenet frame. θ denotes the angle between tangent vectors at stage *s* and *s* + ε, and φ denotes the angle between normal vectors at stage *s* and *s* + ε in the plane of the aperture. They are divided with ε to calculate the normalized curvature and torsion. (**b**) Shape change when parameters are changed in the growing tube model. *C, T*, and *E* represent curvature, torsion, and enlarging ratio, respectively. (**c**) The basic crease pattern for the origami-growing tube model. The black, red, and blue lines show border lines, mountain folds, and valley folds, respectively. The repeated segments are named a “unit” (gray area). The unit height and the left and right angles between the mountain and valley folds are named *D*, α, and β. *D*_*k*_ means the *k*-th unit’s height. The center red line is called “line *l*.” (**d**) An example of the folded shape, in which *D*_*k*_ increases exponentially to mimic a logarithmic spiral (*n* = 10). The numbers on the origami show the unit id. The detail of the folding process is shown in Supplementary Figure 1.

Unlike the above models, the growing tube model’s parameters are biologically meaningful and understandable as the body’s movement relative to the aperture. Moreover, this model suggests that spirals inevitably appear during the organism’s development. The only problem with this model was that it was difficult to assess the parameters from an actual shell shape, but the software reported by Noshita has made parameter estimation possible (Noshita 2014).

The results of these models are usually visualized with computer graphics (Raup 1966; Okamoto 1988). Computer visualization is impressive because it shows the results of calculations more clearly. However, more than visualized data is needed to lead to understanding. In order to understand mathematical models that deal with 3D shapes, it is better to treat the data physically and visually. Origami is one of the cheapest and most easily accessible ways to create 3D structures. So, developing origami-based mathematical models is beneficial for almost everyone, especially students, to understand mathematical models dealing with 3D structures.

In this study, we tried to materialize the growing tube model with origami. There have been some examples of reproducing shell shape forms with origami. Maekawa proposed a flat (planispiral) shell shape origami (Maekawa 2008), which employs the recursion of similar units that mimic the mechanisms of the logarithmic spiral model. Fuse proposed another shell shape origami, employing the recursion of similar units and generating a conispiral shell shape by self-collision (Fuse 2012). These are characterized by the fact that they reproduce a logarithmic helical structure by recursively repeating units of similar shape. However, although they reproduce shell shapes, it is difficult to understand the mathematical model underlying shell shapes.

Therefore, in this paper, we tried to incorporate the growing tube model into origami. We investigated the relationships between the folds connecting repetitive units and the parameters of the growing tube model so that doing origami leads to understanding the model. Furthermore, we showed that various shell morphologies could be reproduced by changing the angle and length of the folds corresponding to the parameters. Based on these data, by changing the torsion periodically, we produced a shape like *Nipponites mirabilis*, one of the famous heteromorph ammonoids. Finally, we investigated the constraints between the folds corresponding to the parameters to clarify whether the origami-growing tube model can reproduce all shell types. This method will be helpful for education in mathematics and biology by disseminating mathematical models.

Results

### Basic crease patterns for the origami-growing tube model

For considering the origami-based growing tube model (origami-growing tube model), we first defined a basic crease pattern for reproducing a shell-like shape. To make the basic crease pattern, we simplified the crease pattern “navel shell,” previously presented by Tomoko Fuse (Fuse 2012). Figure 1c shows an example of the crease pattern. As shown in Figure 1c, we named each part (unit and line *l*), angle (α and β), and length (*D*_*k*_) of the crease pattern. The folding process is shown in Supplementary Figure 1. Figure 1d shows an example of the folded shape that reproduce a planispiral shape (logarithmic spiral). In the following sections, we modified the basic crease pattern partially and observed the folded shape to consider what properties in the crease pattern (e.g., angle) correspond to the parameters in the growing tube model.

### Curvature

We initially investigated what elements in the crease pattern correspond to the curvature in the growing tube model. This section considers the situation of *α* = *β*, and we first considered the situation of *D*_*k*_ = *D* (*constant*). As the corresponding elements, we focused on the unit’s angle α and height *D*. When we varied the angle *α* with a fixed height *D*, the curvature intensity increased as the angle *α* increased (Figure 2a). Next, we changed the height *D* with a fixed angle *α*. In this case, the curvature intensity increased as the height *D* decreased (Figure 2b). Based on these results and the fact that *D* and *α* are independent of each other, the curvature *C* can be estimated as 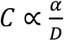. Here, considering the angle between the growth direction of neighboring units after folding (Figure 2c), the curvature can be expressed as 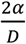, confirming the expression.

**Figure 2:**
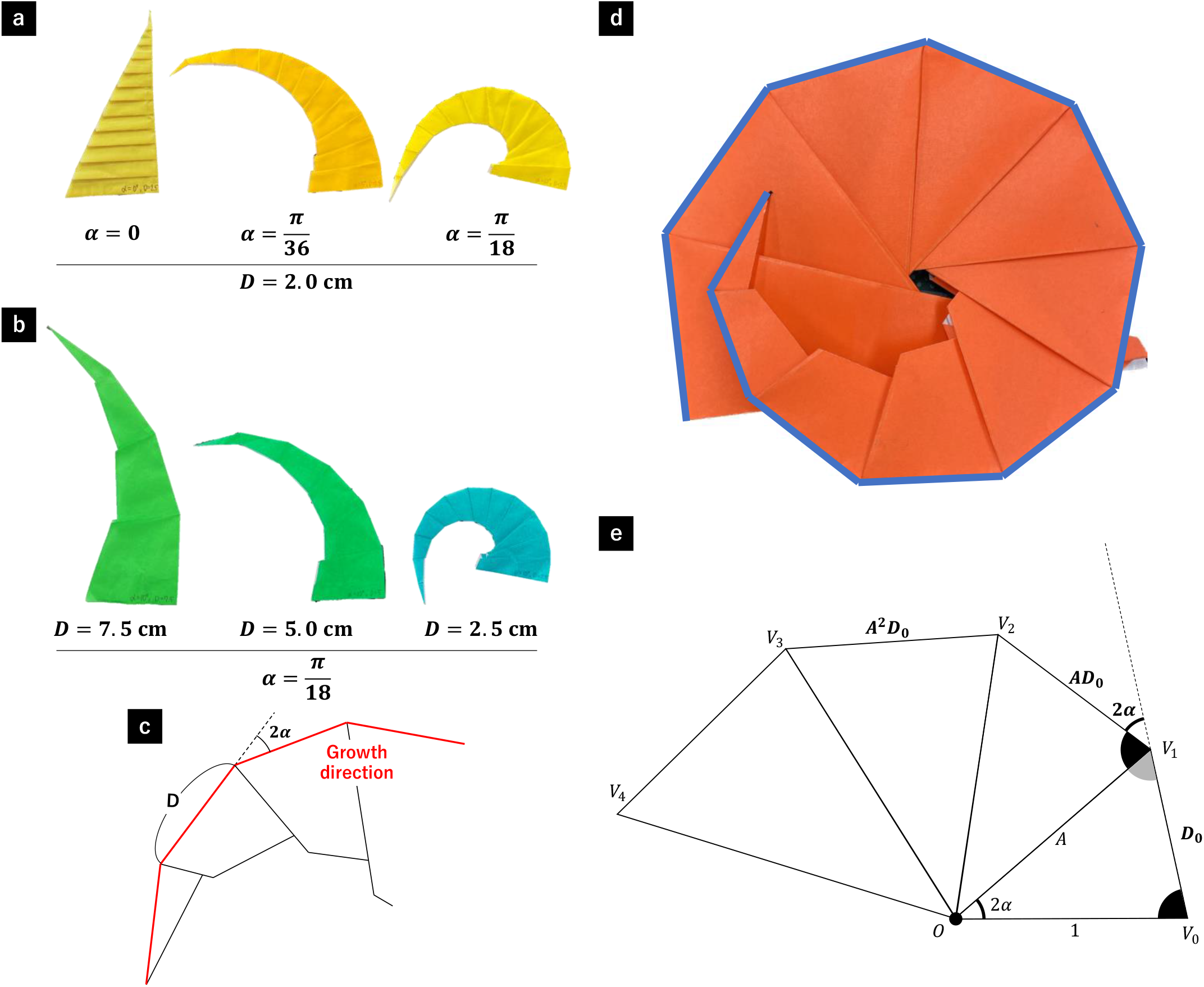
Curvature. (**a-c**) Change in curvature when *D*_*k*_ is constant (= *D*) and *α* is equal to β. (**a**) Change in the curvature when *D* is fixed and only *α* is varied. The values under each figure show the corresponding values of *α* and *D*, respectively. (**b**) Change in curvature when *α* is fixed and only *D* is varied. The values under each figure show the corresponding values of *α* and *D*, respectively. (**c**) The length and angle seen after folding. We named the red line “growth direction,” whose segment length is equal to the unit’s height (= *D*), and the angle between neighboring segments is equal to 2α. (**d**,**e**) Change in curvature when the unit’s height *D*_*k*_ increases exponentially and *α* is equal to β. (**d**) The folded figure when *D*_*k*_ increases exponentially. The spiral of the growth direction approximates a logarithmic spiral. (**e**) The sequence of vertices *V*_0_, *V*_1_, … generated by connected similar triangles have the same trajectory as the growth direction of the folded origami-growing tube model. The parameter of a logarithmic spiral *r* = *B*^θ^ can be calculated from the position of vertices.

Next, we considered the case when *D*_*k*_ is determined by a geometric sequence (*D*_*k*_ = *A*^*k*^*D*_0_), which approximates the logarithmic spiral (Figure 2d). Let us now consider connecting similar triangles in which the ratio of two edges is 1: *A*, and the angle between them is 2*α*, as in Figure 2e (△ *OV*_*i*_*V*_*i*_+1). In this case, the angle between 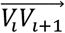 and 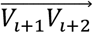 is 2*α* and the ratio of their magnitudes is *A*. So, the sequence of *V*_*i*_ is the same as the trajectory of the growth direction (red line in Figure 2c) of the origami-growing tube model with *D*_*k*_ = *A*^*k*^*D*_0_. When the position of *V*_0_ is set as (1, 0), the position of *V*_*i*_ is (*A*^*i*^ *cos*(2α ∗ *i*), *A*^*i*^ *sin*(2α ∗ *i*)). Because a point on a logarithmic spiral *r* = *B*^θ^ can be expressed as (*B*^θ^ *cos* θ, *B*^θ^ *sin* θ), the parameter *B* of the origami-growing tube model can be expressed as 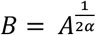.

### Torsion

Next, we examined the crease pattern elements corresponding to the torsion in the growing tube model. The torsion can be understood as the rotation of the body orientation in the plane perpendicular to the growth direction, considering the local coordinate at the aperture (Figure 1a). In the origami-growing tube model, the line *l* in the crease pattern (Figure 1c) or the growth direction in the folded shape (the red line in Figure 2c) can be regarded as a common characteristic point during growth. So, the direction of the normal vector of the origami on the line *l* can represent the organism’s orientation, indicating that the rotation of the normal vector corresponds to the torsion.

As the component corresponding to the torsion, we examined the difference in the angle between the left and right sides (*α* and *β*). Since it is difficult to consider the normal direction using origami alone, we simplified the unit-unit relationship in the origami-growing tube model into the disc-cone model (Figure 3a). In the disc-cone model, we associated the folding process with cutting sectors (2*α* and 2*β*) from a disc to form a cone (Figure 3a, lower lane). Using the model, we considered the trajectory of the line *l* (Figure 3a, red broken lines). In the disc-cone model, the relationship between *n*_0_ and *n*_1_ (the normal vectors of the cone on the lines *V*_0_*V*_1_ and *V*_1_*V*_2_, respectively) is determined by the angles θ and φ (Figure 3b, marked in red). The angle θ is determined by the cone shape, which is determined by the value of *α* + *β*, and can be considered equivalent to the curvature discussed in the previous section. Because φ determines the rotation of the normal vector in the left-right direction, it can be seen as equivalent to the torsion. The value of φ can be calculated as 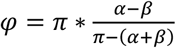.

**Figure 3:**
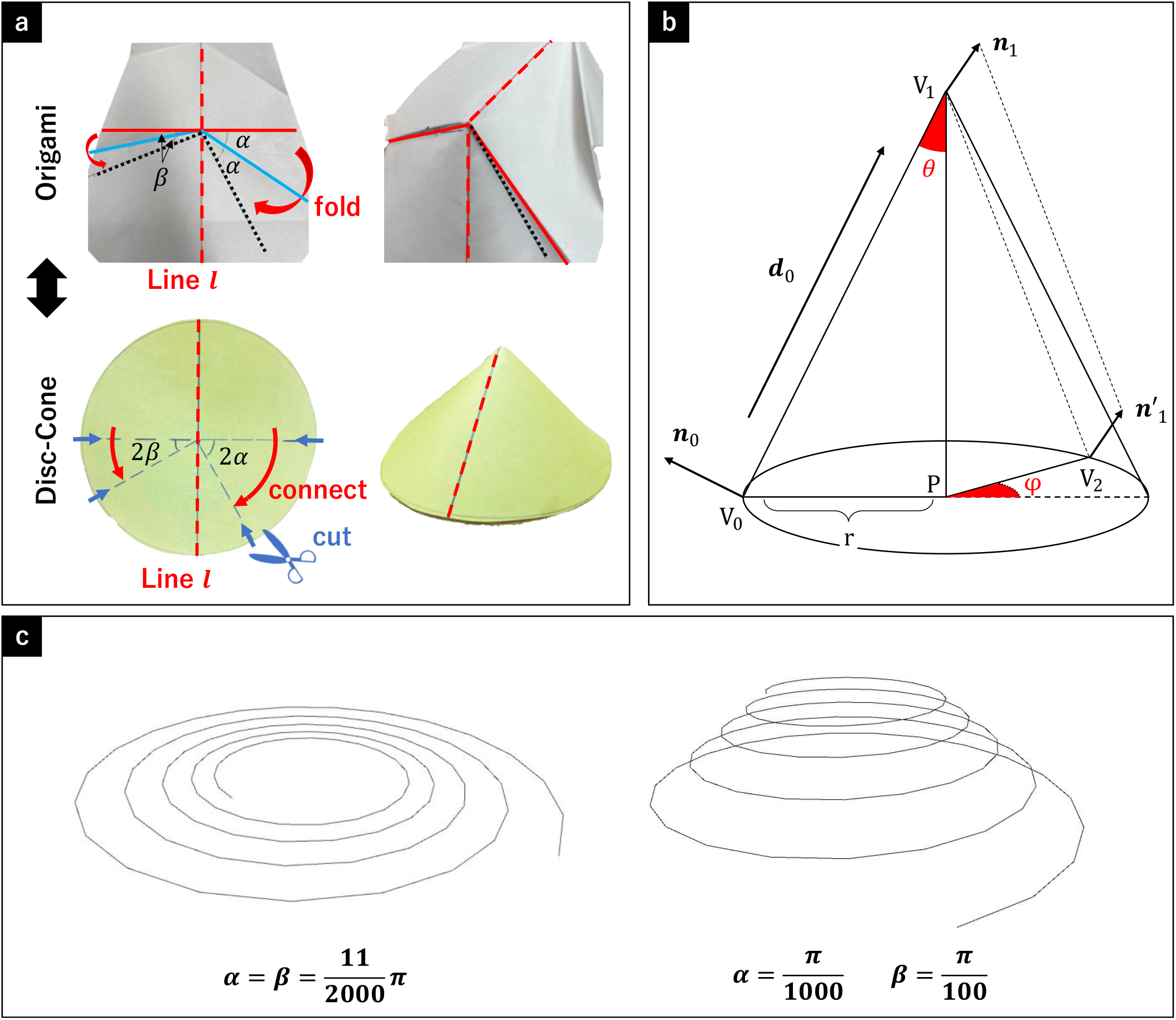
Torsion. (**a**) Association of the unit-unit relationships with the disc-cone model. To simplify origami folding, we regarded the folding process as the process of cutting sectors from a disc to form a cone. The red broken lines show the line *l*. The upper lane shows the origami folding process. The red and blue lines represent the mountain and valley fold shown in Figure 1c. The black dotted lines represent the line to which the red line moves after folding (right image). The lower lane shows the disc-cone model. Cutting the sectors (blue arrows) from the disc and connect the cut lines (red arrows) to form a cone (right image). (**b**) Definition of vectors, vertices, and angles in the disc-cone model, for calculating the next step from the current information. The important angle θ and φ are marked in red. (**c**) Simulation of the line *l* transition based on the disc-cone model (*n* = 99, *A* = 1.01). The left planispiral line shows the result of 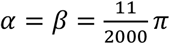. The right conispiral line shows the result of 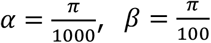.

When *α* is equal to *β* (φ = 0), the result has the same shape as the output generated by the growing tube model with no torsion (Figure 2d). To further confirm the correspondence of φ to the torsion, we conducted computer calculations, in which the relationship between *V*_0_, *V*_1_, *V*_2_, *n*_0_, and *n*_1_ in the disc-cone model were calculated (see supplementary information 1 for the calculation details). We observed the changes when varying the values of *α* and *β*. The results showed that the difference between *α* and *β* regulated the rise in the z-direction (planispiral or conispiral), like the function of the torsion in the original growing tube model (Figure 3c). These findings suggest that φ (or the difference between *α* and *β*) corresponds to the torsion.

### Apex parameters: enlarging ratio and apex sliding

Next, to identify the element corresponding to the radius enlarging ratio, we focused on the angle γ (Figure 4a). We folded the basic crease pattern with various angle γ by altering the height-to-width ratios. Figure 4b illustrates that as the angle γ increased, the enlarging ratio increased (the values of 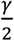 of the dark blue, light blue, and turquoise origami were 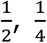, and 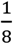, respectively).

**Figure 4:**
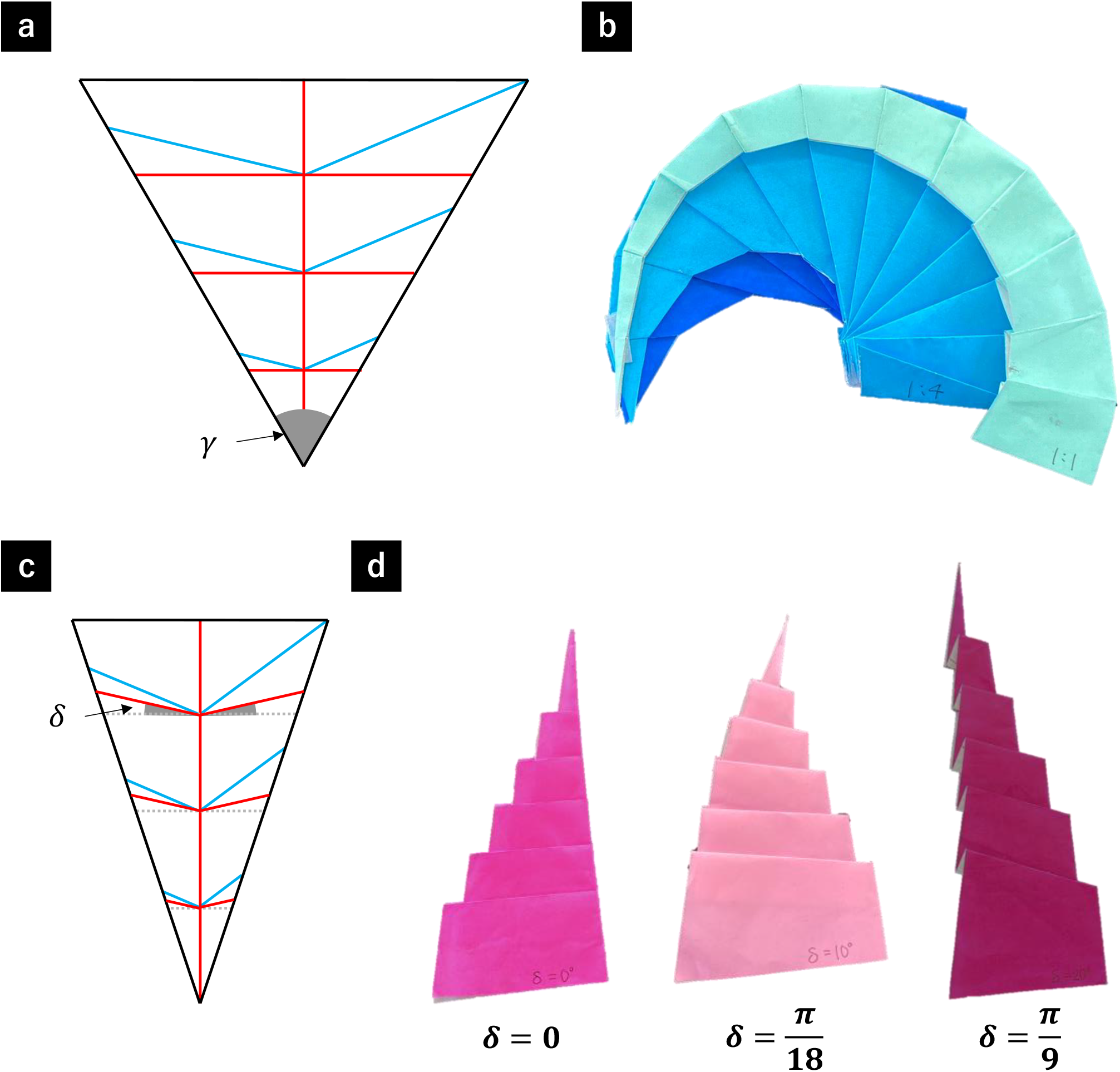
Apex parameters (radius enlarging ratio and apex slide). (**a, b**) The shape change with various height-width ratio. (**a**) We changed the height-width ratio by changing the angle γ. (**b**) The values of 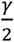 of the dark blue, light blue, and turquoise ones were 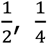 and 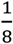, respectively. (**c**,**d**) The shape change with various angle δ. (**c**) We made the parallel mountain folds inclined and named the angle between the mountain folds and the horizontal lines (gray dotted line) angle δ. (**d**) The values of the angle δ of the vivid pink, light pink, and dark pink origami were 0, 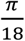, and 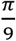, respectively. The apex position slid as the angle δ changed.

In addition, to identify the element responsible for shifting the apex position relative to the aperture, we folded the unit while keeping the angle *α* fixed at 0 and varying the inclination angle *δ* of the unit to 0, 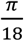, and 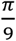 (Figure 4c). As depicted in Figure 4d, as the inclination angle *δ* increased, the magnitude of the apex sliding also increased. This data suggests that the angle *δ* regulate the apex position sliding, which can also be regarded as a change in the orientation of the aperture.

### Reproduction of various shell shapes, including heteromorph ammonoids

Based on the findings, we reproduced various shell shapes using origami (Figure 5 a-d, left images). For comparison, we also reproduced shell shapes with software, in which shapes are generated by the growing tube model (Figure 5 a-d, right images). The similarity between the two sets of shapes demonstrates that the origami-growing tube model effectively replicates the original growing tube model.

**Figure 5:**
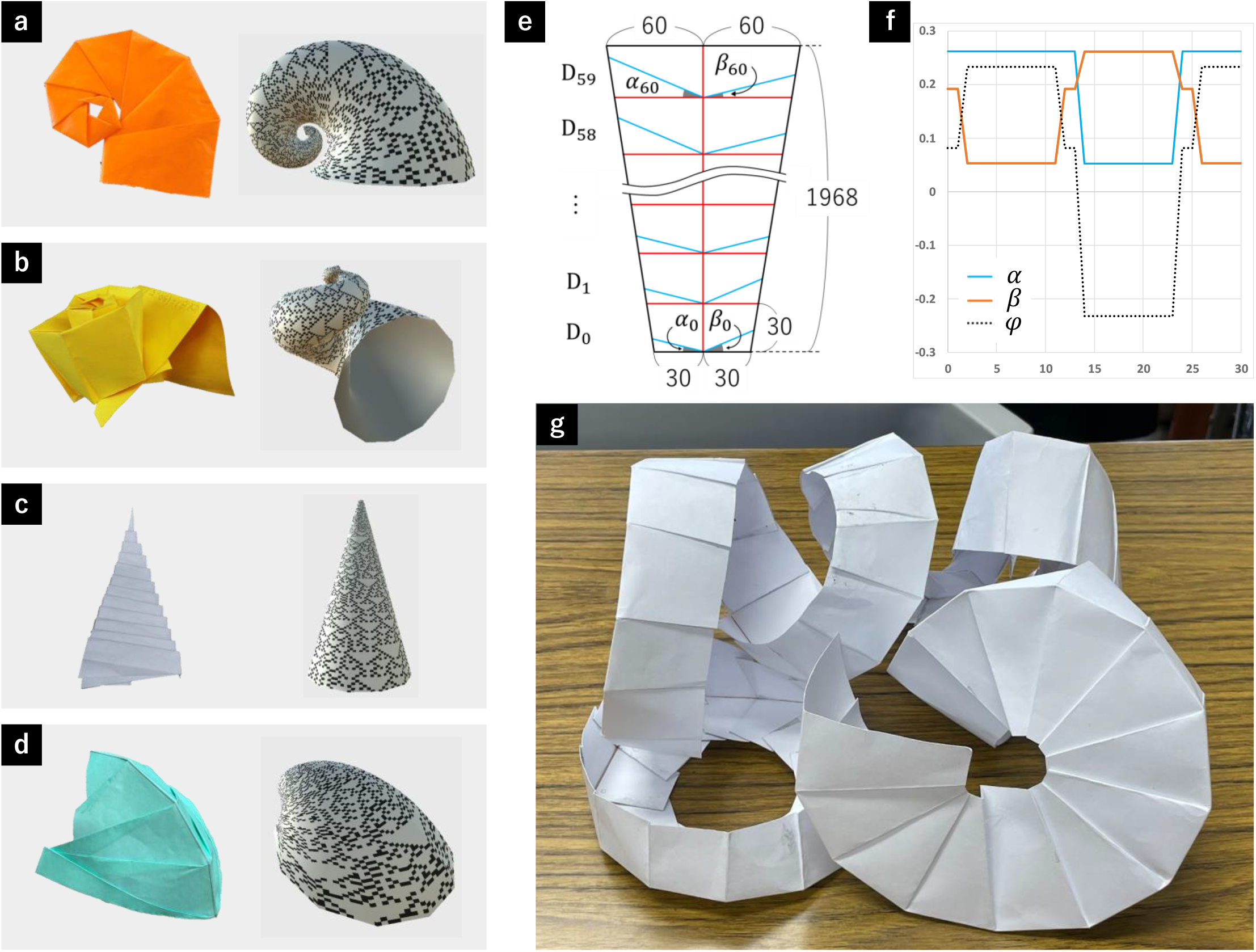
Reproduction of shell shapes. (**a-d**) Various shell shapes reproduced by the origami-growing tube model (left) and the original-growing tube model (right). (**a**) A *Nautilus*-like planispiral shell. (**b**) A gastropod-like conispiral shell. (**c**) An *Orthoceras*-like cone shell. (**d**) A bivalves-like shell. (**e-g**) Reproduction of the heteromorph-ammonoid-like shell shape with origami. (**e**) Crease pattern. We divided the trapezoidal paper into 60 units. The unit of length in the crease pattern is mm. *α* and *β* was changed periodically. (**f**) Diagram of the periodic changes in α, *β*, and φ. The horizontal axis shows the unit number, and the vertical axis shows the magnitude of α, *β* and φ. The unit of angle is radian. (**g**) The folded figure of the crease pattern. Like a heteromorph ammonoid (*Nippoites mirabilis*), The folded figure had a complexly coiled shape.

Furthermore, we attempted to reproduce a heteromorph ammonoid. We folded a trapezoid-shaped paper (Figure 5e). We divided the paper into 60 units, and the unit height increased exponentially (*D*_*k*_ = 1.003^*k*^ ∗ *D*_0_). To mimic the periodic torsion change, we changed (α, *β*) as 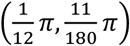, and 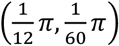 periodically (Figure 5f). As a result, the crease pattern could reproduce a heteromorph-ammonoid-like (*Nipponites mirabilis*-like) shape (Figure 5g).

### Constraints on parameters in the origami-growing tube model

In the origami-growing tube model, the range of reproducibility is limited because of the limitation of the crease patterns (e.g., to prevent the intersection of crease lines). This section considers the situation where *D*_*k*_ = *A*^*k*^*D*_0_ (*A* > 1) and *δ* = 0. First, we considered the relationship between the curvature and torsion. In the origami-growing tube model, the strength of the curvature and torsion can be expressed by (*α* + *β*) and 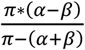. When these indices are denoted with *s* and *t*, the existence conditions of *α* and 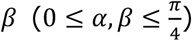 can be written as:

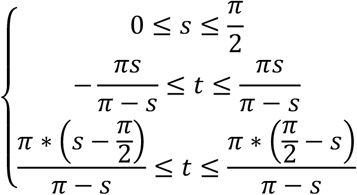

This area is drawn as shown in Figure 6a. These inequalities show that, in the origami-growing tube model, the range in which torsion can be taken is constricted by the value of curvature (Figure 6a). We also considered the relationship between the enlarging ratio and curvature. To simplify the problem, we considered the situation of *α* = *β*. In this situation, by considering the conditions under which blue and red lines do not intersect (as shown in Figure 6b), the following inequality can be derived for the common ratio *A*, angle α, and angle γ.

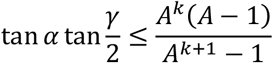

**Figure 6:**
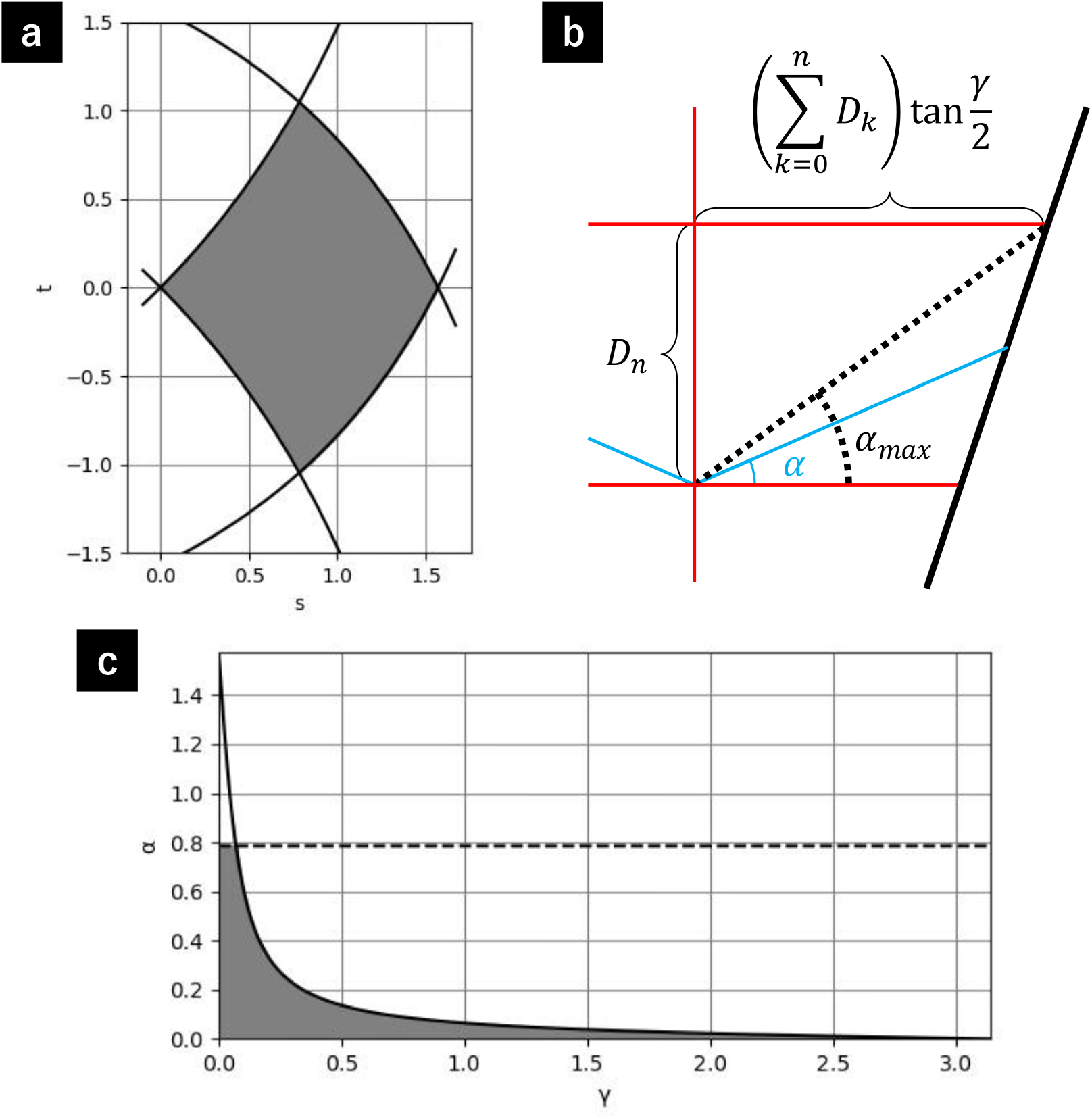
Constraints between parameters in the origami-growing tube model. (**a**) The graph of the relationship between the curvature and torsion. (**b, c**) The range of the angle *α* is constrained to prevent the intersection of crease lines. (**b**) The angle *α* should be more than 0 and no more than α*max* so that the blue line does not intersect with the red horizontal line. (**c**) The graph of the relationship between the enlarging ratio and curvature, under the situation of δ = 0, *α* = *β, A* = 1.03, *nmax* = 59. The dotted line shows 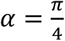 so that the parallel mountain folds do not ride on the line *l*.

The right side of the inequality is a monotonically decreasing function with respect to *k* (Supplementary information 2). So, if the inequality holds for *n* (the maximum value of *k*), it holds for all *k* in 0 ≤ *k* ≤ *n*. Figure 6c shows the relationship between *α* and γ under *A* = 1.03, *n* = 59 (The dotted line shows 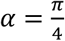 so that the parallel valley folds do not go over the line *l*.). So, the enlarging ratio and curvature constrict each other.

Thus, in the growing tube model, the parameters enlarging ratio, curvature, and torsion, which are independent in the original growing tube model, are not independent. In other words, some shells might not be created with the origami-growing tube model.

## Discussion

In this paper, we have developed an origami-based mathematical model for shell shapes. We identified corresponding folds for each parameter in the growing tube model (curvature, torsion, enlarging ratio). We also found an element for another parameter, apex sliding. Combining these findings, we reproduced various shell shapes, including a heteromorph-ammonoid-like shape. Finally, we investigated the constraints between parameters in the origami-growing tube model.

Previous literature on origami reproduction of shell shapes employs recursively repeating similar units to create logarithmic spirals and generates a conispiral shape by self-collision, not by torsion (Maekawa 2008; Fuse 2012; Lommel 2009). Our origami-growing tube model also employs recursively repeating similar units. However, by introducing the growing tube model into origami, our methods succeeded in generating conispiral shapes by torsion and enabled us to design seashell-like shapes with adjustable parameters. For effective parametric design, understanding the constraints between parameters is crucial. While this study investigated the constraints, further research is needed to fully understand the constraints and expand the range of reproducible shells by reducing the constraints. Compared with the previous literature, our methods have some issues with the similarity of the folded appearance since we focused on reflecting the concept of the growing tube model. For example, the cross-section of the folded structure is not closed. It is necessary to develop more intricate folding techniques while maintaining the clarity of the mathematical model, like curved-line folding in the previous paper (Lommel 2009). Solving these problems may lead to more applications of the origami-growing tube model to design and art.

Mathematical models are usually visualized with computer graphics (Okamoto 1988; Raup 1966), and recent technological advances allow for 3D visualization through naked-eye 3D displays and VR glasses. Our study presents an alternative method for materializing mathematical models through origami, which is universally accessible and cost-effective. So, it is suitable for classroom and home learning environments. Moreover, manually reproducing the results of mathematical models may deepen understanding of the models. Reproducing mathematical models through origami can offer educational benefits as it visualizes and materializes abstract mathematical concepts.

When considering mathematical models that can be better understood using origami, those with fractal or recursive structures seem particularly compatible (shell shapes can also be regarded as recursive structures). Previous research reported that creating the Dragon curve, one of the famous fractal structures, with origami is used for education by combining computer programming (Budinski et al. 2019). Another example of mathematical models with recursive structures is L-systems, which can reproduce plant forms (Prusinkiewicz and Lindenmayer 2012). Incorporating mathematical models appropriately into origami enables more and more people, especially students, to understand the models, leading to future progress in mathematical and theoretical biology. Future research studying origami-based mathematical models is expected.

## Materials and methods

### Origami

We folded the origami (Toyo Corporation, Japan), after manually drawing the crease lines. For creating the heteromorph-ammonoid-like shape, we used a commercially purchased drawing paper (Nippon Notebook Corporation, Japan). The details of how to fold the origami is shown in Supplementary Figure 1.

### Simulation of the torsion

We developed the simulation for visualizing the effect of torsion with processing (Reas and Fry 2007). The calculation details are shown in Supplementary information 1.

## Data Availability

All study data are included in the article and/or Supplementary files.

## Acknowledgements

MU, NF, NY, and YY were supported by Super Science High School (SSH). KM was supported by Grants-in-Aid for JSPS Fellows (DC2), leave a nest grant, and ANRI Fellowship. KM appreciates his colleagues for their helpful comments on this study.

While preparing this work, the authors used DeepL, ChatGPT, and Grammarly to improve the clarity and grammatical usage of English. After using these services, the authors reviewed and edited the content as needed and took full responsibility for the publication’s content.

## References

Budinski, Natalija, Zsolt Lavicza, Kristof Fenyvesi, and Miroslav Novta. 2019. “Mathematical and Coding Lessons Based on Creative Origami Activities.” Open Education Studies 1 (1): 220–27.

Fukutomi, T. 1953. “A General Equation Indicating the Regular Form of Mollusca Shells, and Its Application to Geology, Especially in Paleontology (1).” Bulletin of Geophysics, Hokkaido University 3: 63–82.

Fuse, Tomoko. 2012. Spiral: Origami, Art, Design. Viereck Verlag.

Lommel, A. 2009. “Paper Nautili: A Model for Three-Dimensional Planispiral Growth.” Origami 4. https://doi.org/10.1201/b10653-4/paper-nautili-model-three-dimensional-planispiral-growth-arle-lommel.

Maekawa, Jun. 2008. Genuine Origami: 43 Mathematically-Based Models, from Simple to Complex. Japan Publications Trading.

Moseley, H. 1838. “XVII. On the Geometrical Forms of Turbinated and Discoid Shells.” Philosophical Transactions of the Royal Society of London 128 (0): 351–70.

Noshita, Koji. 2014. “Quantification and Geometric Analysis of Coiling Patterns in Gastropod Shells Based on 3D and 2D Image Data.” Journal of Theoretical Biology 363 (December): 93–104.

Okamoto, T. 1988. “Analysis of Heteromorph Ammonoids by Differential Geometry.” Palaeontology 31 (1): 35–52.

Prusinkiewicz, Przemyslaw, and Aristid Lindenmayer. 2012. The Algorithmic Beauty of Plants. Springer Science & Business Media.

Raup, David M. 1966. “Geometric Analysis of Shell Coiling: General Problems.” Journal of Paleontology 40 (5): 1178–90.

Reas, Casey, and Ben Fry. 2007. Processing: A Programming Handbook for Visual Designers and Artists. MIT Press.

Thompson, Darcy Wentworth, and D’arcy W. Thompson. 1942. On Growth and Form. Vol. 2. Cambridge university press Cambridge.

