## Supplementary Materials for "Origami-based growing tube model for reproducing shell shapes"

Contents

Supplementary Figure 1: How to fold the basic crease pattern.

Supplementary information 1: Disc-cone model.

Supplementary information 2: The constraint condition between $\alpha$ and $\gamma$.

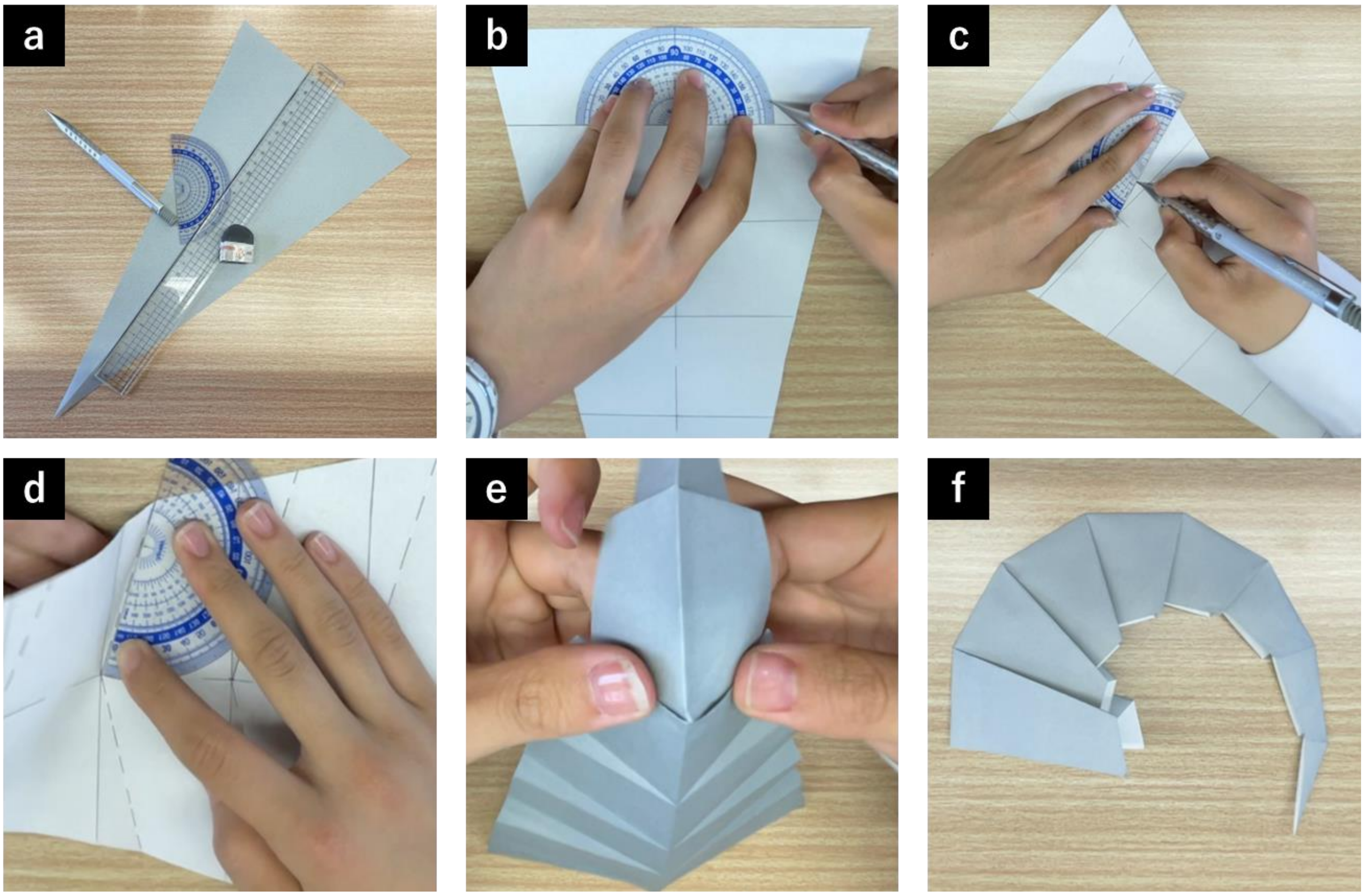


**Supplementary Figure 1: How to fold the basic crease pattern.**

(a) Preparation of tools and origami. We used triangle-shaped origami for the origami-growing tube model (except for heteromorph ammonoids). (b) Drawing lines of mountain folds and marking the angles of $\alpha$ and $\beta$ with a protractor. (c) Drawing lines of valley folds based on the marks. (d) Making the crease based on the lines. (e) Folding the whole origami based on the creases. The overlapping area was pasted with glue in some cases (especially for creating 3D shells). (f) The folded shape.

**Supplementary information 1: Disc-cone model.**

Based on the disc-cone model (Figure 1-1), we calculated the position vectors of $V_2$ and the normal vector $\boldsymbol{n}_1$ (normal vector of the cone on the line $V_1V_2$), using the position vectors of $V_0$ and $V_1$ and the normal vector $\boldsymbol{n}_0$ (normal vector of the cone on the line $V_0V_1$, $\|\boldsymbol{n}_0\| = 1$)).

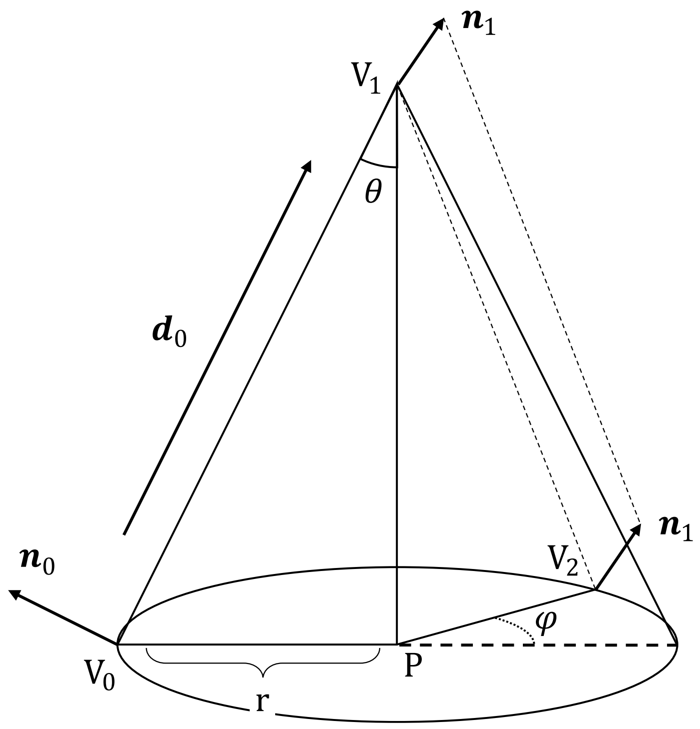


Figure 1-1: the disc-cone model.

First, considering the net figure of the cone (Figure 1-2), the following equation (1) is derived.

$$(2\pi - 2\alpha - 2\beta)\left\|\boldsymbol{d}_0\right\| = 2\pi r$$

$$\therefore \frac{\pi - \alpha - \beta}{\pi} = \frac{r}{\left\|\boldsymbol{d}_0\right\|}(= \sin\theta) \tag{1}$$

Because $\cos\theta$ is more than $0$ when the cone exists,

$$\cos\theta = \sqrt{1 - \left(\frac{r}{\left\|\boldsymbol{d}_0\right\|}\right)^2} \tag{2}$$

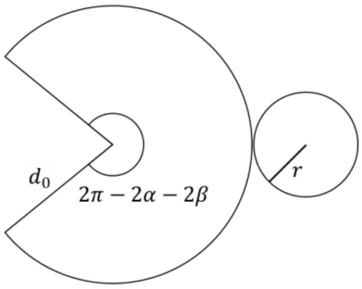


Figure 1-2: the net figure of the cone.

Then, considering the cross-section shown in Figure 1-3, the vector $\overrightarrow{V_0P}$ can be expressed as:

$$\overrightarrow{V_0P} = \left(-d_0 \sin\theta \cos\theta\right)\boldsymbol{n}_0 + \left(d_0 \sin^2\theta\right)\frac{\boldsymbol{d}_0}{d_0} = \left(-d_0 \sin\theta \cos\theta\right)\boldsymbol{n}_0 + \left(\sin^2\theta\right)\boldsymbol{d}_0 \quad (3)$$

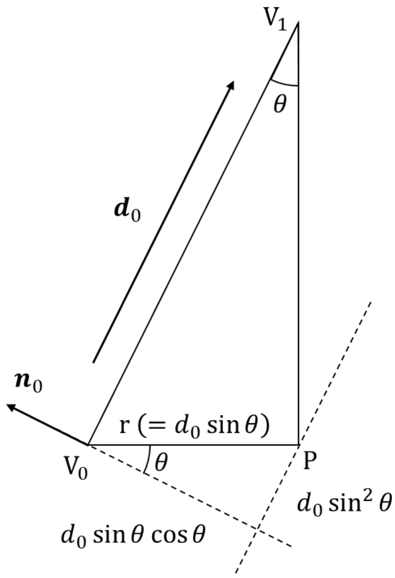


Figure 1-3: the cross-section of the cone at the point $V_0$ and $P$.

Here, considering the base of the cone (Figure 1-4), the position of $V_2$ can be expressed as:

$$\overrightarrow{PV_2} = (\cos\varphi)\overrightarrow{V_0P} + \left(d_0 \sin\theta \sin\varphi\right)\frac{\boldsymbol{n}_0 \times \boldsymbol{d}_0}{d_0} = (\cos\varphi)\overrightarrow{V_0P} + (\sin\theta \sin\varphi)\boldsymbol{n}_0 \times \boldsymbol{d}_0 \quad (4)$$

, where $\cos\varphi$ and $\sin\varphi$ are calculated numerically on a computer with the value of $\varphi = \frac{\pi*(\alpha-\beta)}{\pi-\alpha-\beta}$.

Using the equations $(1-4)$, the position of $V_2$ can be calculated.

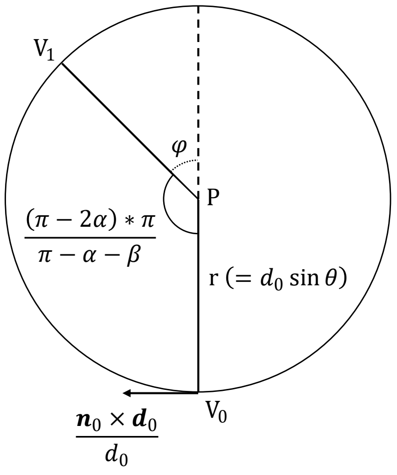


Figure 1-4: the base of the cone.

Regarding the normal vectors, $\boldsymbol{n}_1$ is the result of rotating $\boldsymbol{n}_0$ around $\overrightarrow{PV_1}$ counterclockwise by an angle $(\pi + \varphi)$. So, $\boldsymbol{n}_1$ can be calculated as:

$$\boldsymbol{n}_1 = \big(1 - \cos(\pi + \varphi)\big) \left( \boldsymbol{n}_0 \cdot \frac{\overrightarrow{PV_1}}{\left\| \overrightarrow{PV_1} \right\|} \right) \frac{\overrightarrow{PV_1}}{\left\| \overrightarrow{PV_1} \right\|} + \boldsymbol{n}_0 \cos(\pi + \varphi) - \left\{ \frac{\overrightarrow{PV_1}}{\left\| \overrightarrow{PV_1} \right\|} \times \boldsymbol{n}_0 \right\} \sin(\pi + \varphi)$$

$$= \big(1 + \cos\varphi\big) \left( \boldsymbol{n}_0 \cdot \frac{\overrightarrow{PV_1}}{\left\| \overrightarrow{PV_1} \right\|} \right) \frac{\overrightarrow{PV_1}}{\left\| \overrightarrow{PV_1} \right\|} - \boldsymbol{n}_0 \cos\varphi + \left\{ \frac{\overrightarrow{PV_1}}{\left\| \overrightarrow{PV_1} \right\|} \times \boldsymbol{n}_0 \right\} \sin\varphi$$

So, using $\overrightarrow{PV_1} = (d_0 \sin\theta \cos\theta)\boldsymbol{n}_0 + (1 - \sin^2\theta)\boldsymbol{d}_0$, it is possible to calculate $\boldsymbol{n}_1$ numerically on a computer.

**Supplementary information 2: The constraint condition between α and γ.**

We considered the following inequality.

$$\tan\alpha \tan\frac{\gamma}{2} \leq \frac{A^k(A-1)}{A^{k+1}-1} \quad (A > 1.0, k \geq 0)$$

To consider the characteristics of the right-side of the inequality, we named it as $f(A,k)$.

$$f(A,k) = \frac{A^k(A-1)}{A^{k+1}-1} = \frac{A-1}{A-\frac{1}{A^k}}$$

Then, we compared $f(A,k)$ with $f(A,k+1)$. Because $f(A,k) > 0$,

$$f(A,k) > f(A,k+1) \Leftrightarrow \frac{f(A,k)}{f(A,k+1)} > 1 \Leftrightarrow \frac{A-\frac{1}{A^{k+1}}}{A-\frac{1}{A^k}} > 1 \Leftrightarrow A-\frac{1}{A^{k+1}} > A-\frac{1}{A^k} \Leftrightarrow \frac{1}{A^{k+1}} < \frac{1}{A^k}$$

So, $f(A,k)$ is a monotonically decreasing function with respect to $k$.
